# Proteogenomic Profiling Reveals a Distinct Endogenous p16INK4a-Associated Senescence Signature in the Human Ovary

**DOI:** 10.64898/2026.02.02.703302

**Authors:** Fiona Senchyna, Kevin Schneider, Pooja Raj Devrukhkar, Nicolas Martin, Tommy Tran, Josef Byrne, Mark A. Watson, Fei Wu, Minja Belic, Matias Fuentealba, Elisheva D. Shanes, Mary Ellen G. Pavone, Bikem Soygur, Birgit Schilling, Simon Melov, Francesca Duncan, David Furman

## Abstract

The tumor suppressor and cell cycle regulator, p16^INK4a^ (p16), has been extensively linked to cellular senescence, and its accumulation can reflect endogenous senescence within ovarian tissue. However, gene and protein signatures associated with p16 have not been well defined in human tissue. We utilized immunohistochemical (IHC) staining for P16 to identify distinct positive (P16+) and negative (P16-) regions within the ovarian cortex and employed the GeoMx Digital Spatial Profiler for simultaneous proteomic and transcriptomic analyses on cortical tissue cores. Differential expression and translation between p16-positive and p16-negative cores identified genes and proteins that are cellular senescence related (e.g., *CDKN1A, GADD45B, GADD45G*, and *MYC*) or key regulators of the extracellular matrix (*e*.*g*., collagen I, *ADAMTS4*, and *MMP11*). Additionally, the transcriptomic signature identified here was significantly enriched for the spatially derived ovarian p16-associated signature, BuckSenOvary, but not for other senescence gene sets. Lastly, given the association between changes to the extracellular matrix in aged ovaries and ovarian cancer, we compared genes upregulated and downregulated in p16-positive regions relative to p16-negative regions against multiple ovarian cancer transcriptomic datasets. These findings provide new insight into the molecular landscape of naturally occurring ovarian senescence and its possible relationship to age-associated disease processes, including cancer development.

## Introduction

Cellular senescence is a fundamental hallmark of aging and has been increasingly implicated in ovarian aging and functional decline [1, 2]. Senescence-related pathways are enriched in aged ovaries compared to younger donors [3]. In a non-human primate ovary spatial transcriptomic atlas, the proportion of senescent cell populations increased with age [4], and in mice, the upregulation of senescence-associated markers precede depletion of the ovarian reserve [5]. In post-menopausal women, cessation of ovarian function following depletion of ovarian reserve in the cortex is associated with worsened health outcomes and increased mortality risk [6].

Recent efforts to define and quantify cellular senescence in human tissue have stimulated a wealth of spatial profiling studies which are now producing a great deal of novel biomarkers and cell senescence profiles (or “senotypes”) [7]. Yet, senotypes based on multiple tissue sources and cell types often do not include the ovary [8,9].

p16^INK4a^ (p16) is a tumor suppressor and key regulator of the cell cycle that serves as a well-established marker for cellular senescence [10-13]. Here, we investigate whether distinct areas of the post-menopausal ovary exhibit differences in cellular senescence, by utilizing the GeoMx Digital Spatial Profiler (DSP). We performed simultaneous proteomic and transcriptomic profiling of ovarian cortical tissue cores classified as p16-positive or p16-negative by p16^INK4a^ immunoreactivity in three post-menopausal women. Parallel transcriptomic and proteomic analysis allowed us to examine the overlap in signatures, which are not always aligned due to post-transcriptional regulation and protein turnover dynamics [14]. Differentially expressed genes in p16-positive relative to p16-negative regions were assessed for enrichment of a recently identified ovarian p16-associated signature, BuckSenOvary [15], along with other senescent datasets. Finally, due to the association between ovarian aging, fibrosis, cellular senescence and tumorigenesis [16, 17], we further examined the enrichment of the p16 and doxorubicin ovarian senescence signatures across multiple ovarian cancer datasets.

## Results

All participants were Caucasian and post-menopausal (aged 52, 74, and 74) with BMIs in the healthy to overweight range and confirmed to be free of any ovarian pre-malignancy or malignancy. Two cores representing an identified p16-positive and p16-negative region within the ovarian cortex were taken for each participant (Figure 1A). Transcriptomic and proteomic profiling was performed on regions of interest (ROIs) spanning the core, with 8–12 ROIs per core. Dimensional reduction using UMAP showed clustering of ROIs first by core and then by participant (Figure 1B). Participant 2 and Participant 3 clustered together, distinct from Participant 1, suggesting that participant age may underlie this separation given the 22-year difference. Cell type deconvolution of pseudobulk profiles identified six cell types (Figure 1C): Stroma 1, Stroma 2, perivascular cells, endothelial cells, smooth muscle cells, and epithelial cells. Consistent with the published single-cell dataset used as the deconvolution reference [18], Stroma 1 and Stroma 2 constituted the majority of cells, though they comprised a larger proportion in the tissue cores than in the original single-cell data (>90% between Stroma 1 and Stroma 2 in each ROI as compared to approximately 50% in the single cell). Based on prior characterization, Stroma 1 and Stroma 2 represent biologically distinct populations enriched for lipid metabolism and cell-matrix adhesion functions, and matrix fibroblast signatures, respectively. Notably, the only cell type absent from the single cell reference were immune cells. This could be due to a combination of low abundance of immune cells in the cores, limitations of the reference which contained only three samples, and high diversity of immune cell subtypes. Lastly, p16-positive and p16-negative ROIs and aggregated cores did not cluster by cell type, indicating the p16-associated signature may be independent of cell composition.

**Figure 1.**
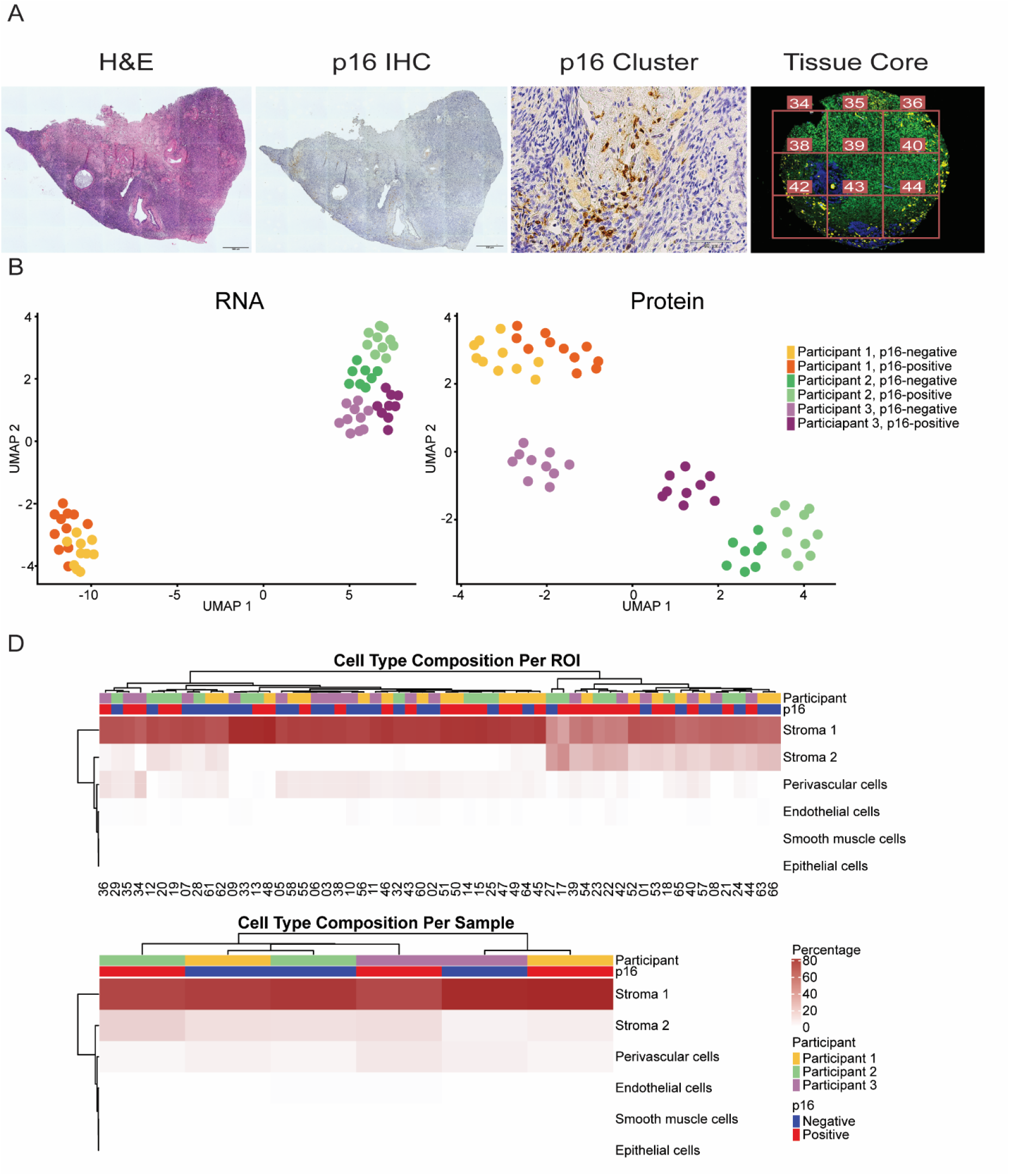
Identification and comparison of p16 positive and negative regions in the ovarian cortex. (A) Representative tissue from a participant showing morphology from H&E image and p16 immunostaining (yellow). Regions of interest (ROIs), numbered and covering the full area of the core, were subject to simultaneous proteomic and transcriptomic profiling. (B) UMAP plot of the ROIs in the RNA and protein datasets, respectively, show clustering to the core and then the participant level, with Participant 2 and Participant 3 clustering separately from Participant 1, likely due to age differences between the participants. (D) Heatmaps of the cell type deconvolution reveal most cells in each ROI and aggregated core are stromal 1 and stromal 2 cells, indicating differences between p16-positive and p16-negative regions are not based on cell type composition.

To assess biological differences between p16-positive and p16-negative regions in the post-menopausal ovarian cortex, we performed differential analysis using both whole-transcriptome profiling and a 570-plex proteomic panel. In p16-positive regions relative to p16-negative regions. 126 transcripts and 7 proteins were upregulated, while 81 transcripts and 1 protein were downregulated (Figure 2A; Supplementary Tables 1–2). The mean correlation between transcript and protein abundance across all analyzed genes was 0.03 (Figure 2B). Directional agreement was observed across several genes that were significantly differentially expressed in either ‘omics and found in both (Figure 2C). A notable exception was *MYC*, a transcription factor and well-established proto-oncogene that regulates a wide range of processes such as cell cycle progression and metabolism. Dysregulation and overexpression of *MYC* is observed in many forms of cancer and is known to suppress senescence [19]. However, it has also been implicated in oncogene-induced senescence (OIS) through transcriptional activation of p14ARF and subsequent stabilization of p53 [20]. In the present study, *MYC* was significantly upregulated at the transcript level (*p*-value = 3.57 × 10^-5^) but downregulated at the protein level. This discordance between mRNA and protein levels suggests post-transcriptional or post-translational regulation of *MYC* during senescence and/or by the autoregulatory negative feedback loop of *MYC*, wherein excessive *MYC* activity triggers transcriptional repression or protein degradation to limit its own accumulation [21].

**Figure 2.**
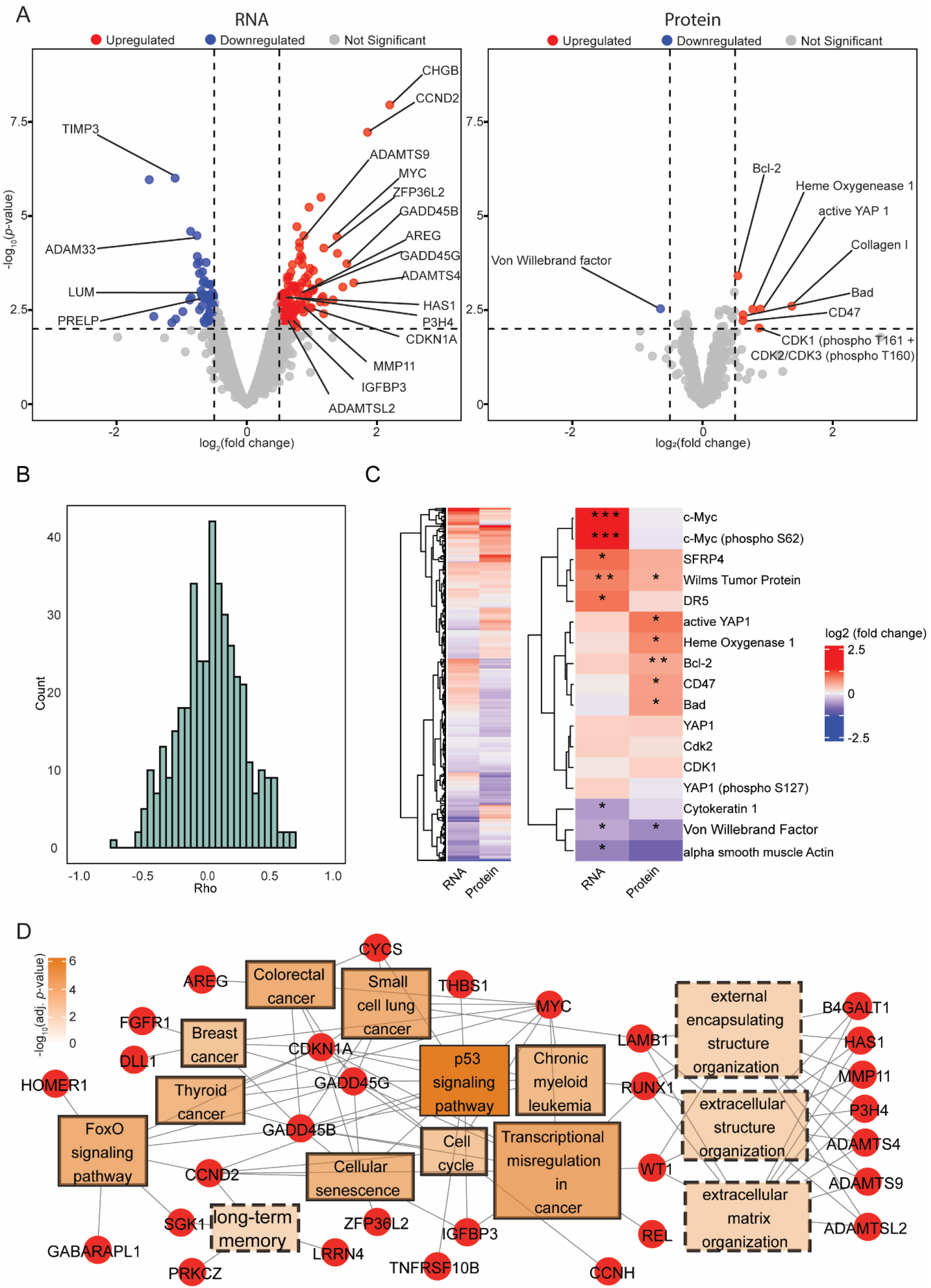
Identification and pathway analysis of p16-positive relative to p16-negative differentially expressed genes and translated proteins. (A) Volcano plots of the 126 upregulated and 81 downregulated transcripts and 8 upregulated and 1 downregulated proteins associated with differential expression and translation between p16-positive and p16-negative regions. (B) The spearman correlation coefficients between transcripts and proteins of the same gene (*n*=417) in the same ROI varied greatly depending on the gene. (C) A heatmap of the common genes and proteins that were differentially expressed and differentially translated, respectively, reveal modest agreement in directionality. However, many genes that were significantly changing in either the protein or RNA dataset were directionally aligned (logFoldChange >0.5, *p*-val * = < 0.01, ** = < 0.001, *** = < 0.0001). (D) Network showing the p16-positive upregulated genes (ellipse) and their association with significantly enriched GO biological processes (rectangle with dashed line) and KEGG (rectangle with solid line) pathways, including multiple extracellular matrix and cellular senescence associated pathways.

### p16 signature is related to cellular senescence and changes in the ECM

Pathway overrepresentation analysis of significantly upregulated genes identified several KEGG pathways associated with cellular stress response and growth arrest, including cellular senescence, p53 signaling, and FoxO signaling pathways. Both the p53 and FoxO pathways are known to be activated by diverse cellular stressors such as oxidative stress, oncogenic signaling, DNA damage, and telomere shortening [22]. Activation of these pathways initiates anti-proliferative responses and subsequently tumor suppressive mechanisms, including apoptosis and cellular senescence [23] (Figure 2D). Several notable genes were associated with these pathways including *CDKN1A* (p21), a cyclin-dependent kinase inhibitor and canonical marker of senescence that is transcriptionally regulated by p53 and FoxO signaling to induce cellular senescence [24, 25], and *GADD45B* and *GADD45G*, members of the Growth Arrest and DNA Damage-inducible 45 family, that are known to mediate growth arrest and DNA repair in response to genotoxic stress [26]. In the proteomics analyses, Bcl-2 and Bad, members of the Bcl-2 family proteins, were upregulated. Bcl-2 is an anti-apoptotic protein associated with cell survival of senescent cells, while Bad, in response to stress, is known to be upregulated and bind to Bcl-2 to sequester its anti-apoptotic effects [27-29].

Interestingly, *CCND2*, which encodes cyclin D2, a regulator that promotes G1-to-S phase transition and cellular proliferation, was also enriched in p16-positive regions [30]. This finding may reflect a compensatory or context-dependent response to senescence-associated proliferative arrest, as cyclin D2 has been shown in some systems to participate in early stress signaling prior to full cell cycle exit [31].

Gene ontology analysis revealed significant enrichment of extracellular matrix (ECM)-related pathways in the p16-positive regions relative to p16-negative regions (Figure 2D). These findings were supported in the proteomics analysis by the identification of increased levels of collagen I, a protein linked to ovarian fibrosis, aging, and cellular senescence [32], and active YAP1, a mechanosensitive transcriptional regulator that has been previously linked to a stiffened cortex and reproductive aging [33]. Several genes associated with collagen assembly, including *LUM, P3H4*, and *PRELP*, were differentially expressed; downregulation of *LUM* has previously been associated with ECM instability and dysregulated collagen organization [34]. ECM-associated changes included differential regulation of multiple matrix-remodeling enzymes such as *MMP11, TIMP3, ADAM33, ADAMTS9, ADAMTSL2, ADAMTS4*, and *HAS1* [35, 36].

### Transcriptomic signature shows significant enrichment for the p16 associated signature, BuckSenOvary, but weak enrichment of other known senescence gene sets

To investigate whether the p16-associated profile found here was concordant with other known senescence gene sets, we performed gene set enrichment of the differentially expressed genes identified here against several senescence gene sets, including a recently derived spatially resolved p16-associated transcriptomic signature (BuckSenOvary) [15], a set of upregulated genes an in-vitro doxorubicin-induced senescence ovarian cortex model [18], SenMayo [8], the SenePy universal signature [9], GO cellular senescence, and Reactome cellular senescence. While all senescence gene sets exhibited positive normalized enrichment scores, only BuckSenOvary reached statistical significance (adjusted *p*-value < 0.05; Figure 3A). We observed partial concordance between the 32 genes in BuckSenOvary and our transcriptional profile, with two genes significantly upregulated (*CCND2* and *MYC*) and one significantly downregulated (*TXNIP*) in our dataset (Figure 3B). This partial overlap likely reflects differences in donor populations and transcriptional profiling over tissue cores rather than manually selected regions. None of the genes in BuckSenOvary were significantly upregulated in the p16-positive regions compared to the p16-negative regions in the proteomics analysis. However, this analysis was limited as only 5 genes from BuckSenOvary were present in the proeome panel (Figure 3B).

**Figure 3.**
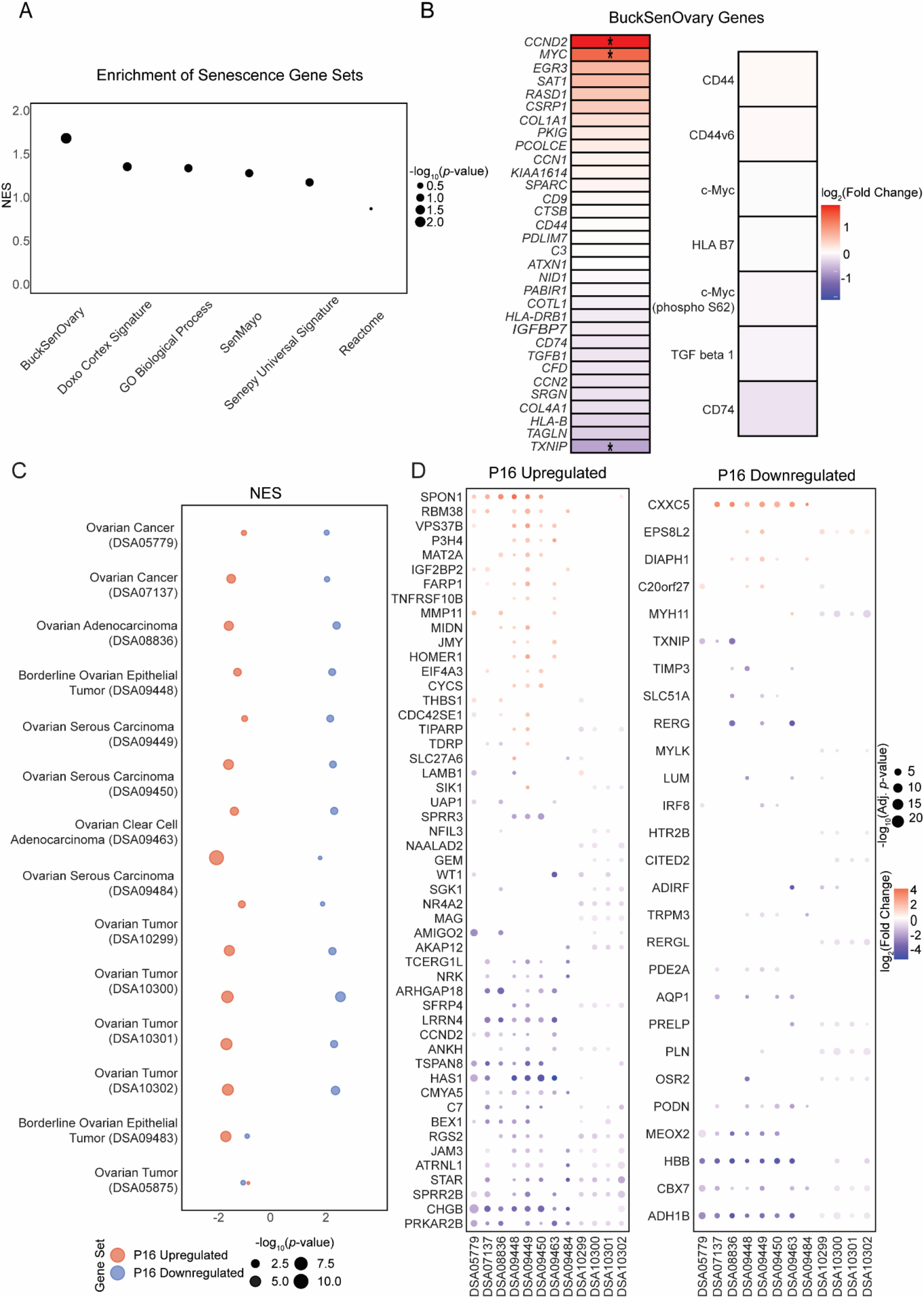
Comparison of the p16-associated cortex profile with other senescence gene sets and enrichment of p16-positive upregulated and downregulated genes in ovarian cancer datasets. (A) The p16-associated profile derived from the ovarian cortex here was significantly positively enriched for BuckSenOvary (normalised enrichemnt score=1.68, adjusted *p*-value<0.05). (B) Differential expression of the individual gene set in BuckSenOvary reveals not all genes are in directional agreement, likely due to differences in individual participants and p16-positive region selection methodology. None of the limited number of BuckSenOvary genes in the proteome panel were significantly differentially expressed. (C) Genes upregulated in the p16-positive regions were negatively enriched in ovarian cancer signatures from DiSignAtlas, while many of the datasets were positively enriched for the p16-positive downregulated genes, meaning many genes were downregulated in both p16-positive regions and cancer. (D) dot plots of genes from the p16 gene sets that are also significantly changed in cancer datasets, including those related to the ECM (e.g., *TIMP3, PODN, LUM*) (*p*-adj < 0.01 and −0.5>log fold change<0.5).

### Many ECM and cell cycle related genes in p16 signature are also dysregulated in cancer

To examine the relationship between senescence-associated signatures and ovarian cancer, gene set enrichment of the p16-associated upregulated and downregulated genes was applied to ovarian cancer datasets from the DiSignAtlas [37] repository. 30 datasets were labeled as ovarian cancer, tumor, or carcinoma from ovary tissue, however, upon further inspection 15/30 were excluded due to being either benign tumors (*n*=3), tissue not from site of tumor (*n*=3), cell lines (*n*=7), small RNA sequencing (*n*=1), or dataset with no associated publication (*n*=1) (list of excluded datasets in Supplementary Table 3). The remaining 15 datasets consisted of genes that were differentially expressed between ovarian tumor tissue and control samples, described as normal ovarian surface epithelia or normal ovary samples. They encompassed a range of disease stages and histopathological subtypes, including high-grade serous carcinoma, borderline ovarian tumors, and clear cell ovarian cancer (Table 1).

**Table 1.** Enrichment of p16-positive associated upregulated and downregulated genes in cancer datasets. Normalized enrichment score and significance of p16-positive associated upregulated and downregulated genes in various ovarian cancer datasets. Gene set enrichment score calculated with fgsea. DSAID=Disign Atals ID.

| DSaID | Pubmed ID | Case Samples Description (N) | Control Samples Description (N) | Normalized Enrichment Score (adjusted *p*-value) |  |
| --- | --- | --- | --- | --- | --- |
|  |  |  |  | P16 Up | P16 Down |
| DSA05779 | 18593951, 25944803 | Primary ovarian tumors from untreated late-stage (III-IV) ovarian cancer patients (185) | normal ovarian surface epithelium (10) | -1.3 (1.21E-1) | 1.3 (1.26E-1) |
| DSA05875 | 29321820 | Ovary tumor from patients with stage II or III, or grade 3 serous carcinoma (8) | normal ovarian surface epithelium (5) | -1.2 (2.45E-1) | -1.3 (1.61E-1) |
| DSA07137 | 28199976 | Ovarian epithelial tumor samples from patients with high grade serous ovarian cancer (16) | normal ovarian surface epithelium (6) | -1.7 (**1.25E-3**) | -1.38 (1E-1) |
| DSA08836 | 20040092 | Ovarian surface epithelia from post-menopausal women with serous adenocarcinoma (12) | normal ovarian surface epithelium from peri- and post-menopausal patients (12) | -1.83 (**5.36E-4**) | 1.4 (1.36E-2) |
| DSA09448 | 21451362 | Paraffin-embedded sections of tumors from serous borderline ovarian tumors (8) | normal ovarian surface epithelium (6) | -1.5 (**8.14E-3**) | 1.5 (**1.88E-2**) |
| DSA09449 |  | Paraffin-embedded sections of tumors from low-grade serous ovarian carcinomas (13) | normal ovarian surface epithelium (6) | -1.28 (6.20E-2) | 1.45 (**1.97E-2**) |
| DSA09450 |  | Paraffin-embedded sections of tumors from high-grade serous ovarian carcinomas (22) | normal ovarian surface epithelium (6) | -1.8 (**1.16E-4**) | 1.54 (**2.09E-2**) |
| DSA09463 | 21754983 | clear cell ovarian cancer specimens from primary tumors of untreated ovarian cancer patients (10) | normal ovarian surface epithelium (10) | -1.6 (**3.05E-3**) | 1.6 (**1.4E-2**) |
| DSA09483 | 23029477 | serous ovarian borderline tumor (4 from 8 samples pooled based on histological classification and stage) (8) | normal ovarian surface epithelium (4) | -1.9 (**4.94E-5**) | 1.6 (1.76E-1) |
| DSA09484 |  | Serous ovarian carcinoma (4 from 11 samples pooled based on histological classification and stage) | normal ovarian surface epithelium (4) | -2.1 (**9.04E-10**) | 1.1 (2.34E-1) |
| DSA09527 | 24666724 | ovarian tumor from serous patients with papillary adenocarcinoma (10) | normal ovary samples (10) | -1.4 (**1.88E-2**) | 1.2 (1.83E-1) |
| DSA10299 | 16452189, 19843521, 17418409, 27538791, 18945643 | Ovarian tumor from patients with clear cell carcinomas (8) | normal ovary samples (4) | -1.8 (**6.29E-5**) | 1.5 (**1.4E-2**) |
| DSA10300 |  | Ovarian tumor from patients with ovarian endometrioid adenocarcinoma (37) | normal ovary samples (4) | -1.8 (**3.42E-7**) | 1.8 (**1.18E-4**) |
| DSA10301 |  | Ovarian tumor from patients with mucinous ovarian carcinomas (13) | normal ovary samples (4) | -1.9 (**7.54E-6**) | 1.6 (**1.4E-2**) |
| DSA10302 |  | Ovarian tumor from patients with serous ovarian carcinomas (41) | normal ovary samples (4) | -1.8 (**7.76E-6**) | 1.6 (**2.18E-3**) |

Overall, the p16-associated upregulated genes were negatively enriched, while the p16-downregulated genes were positively enriched in the cancer datasets, meaning many genes that were downregulated in p16 were also downregulated in the ovarian cancer datasets (Figure 3C, Table 1). Of those genes that were significantly downregulated in p16-positive regions relative to p16-negative regions, 22 genes were significantly downregulated in at least 3 cancer datasets and significantly upregulated in none (*ADH1B, CBX7, HBB, MEOX2, PODN, AQP1, OSR2, PLN, PRELP, PDE2A, RERG, RERGL, SLC51A, TIMP3, TRPM3, TXNIP, ADIRF, CITED2, HTR2B, IRF8, LUM*, and *MYLK*) (Figure 3D displays those p16 genes found in at least 2 cancer datasets, the full list can be found in Supplementary Table 4). Many are extracellular matrix (ECM)-related genes, including *TIMP3, PODN, LUM*, and *PRELP*. These findings are consistent with the established role of fibrosis and ECM remodeling in shaping the tumor microenvironment during cancer progression [16, 34], although the relevance of these changes to ovarian tumorigenesis warrant further investigation given that many ovarian cancers originate in the fallopian tube and disseminate to sites such as the omentum and parietal peritoneum [38, 39].

Beyond ECM remodeling, several genes associated with cell cycle regulation (*CCND2, SGK1, WT1*), apoptosis (*SFRP4, AKAP12, WT1, SIK1*), and cellular senescence (*CCND2, MYC, IGFBP3*) display opposing expression patterns between the p16 signature and cancer. The most significantly upregulated gene in the p16 signature, *CHGB*, which encodes Chromogranin B, a protein involved in catecholamine secretion, was significantly downregulated in most cancer signatures (8/15). Decreased *CHGB* expression has previously been linked to chronic stress-induced breast cancer [40].

## Discussion

Ovarian aging is a critical biological process with far-reaching consequences for both fertility and overall health. Cellular senescence is recognized as a central hallmark of this process and fully elucidating how senescence manifests within the ovary and influences its surrounding microenvironment remains an active area of investigation [1]. In this study, we validated a previously defined p16-associated transcriptional signature, BuckSenOvary [15], in independent human donors and integrated proteomic profiling to characterize ovarian senescence. The little concordance between transcriptomic and proteomic abundance of the same genes was not unexpected, given known influences including differences in translation efficiency and the shorter half-life of mRNA relative to proteins [11]. Despite this, many of the differentially expressed genes and translated proteins were linked to the same biological processes. For instance, upregulation of Collagen I and active YAP I at the proteomic level and changes to ECM related genes (e.g., *MMP11, TIMP3, ADAM33*, and *ADAMTS4*) in the transcriptomics are consistent with fibrosis [15]. Similarly, differential expression of genes such as *CDKN1A, GADD45B*, and *GADD45G* and differential translation of the proteins Bad and Bcl-2 suggest involvement of the p53 signaling pathway and cellular senescence.

While spatial selection of p16-positive regions for transcriptional analysis is not always practical, tissue core-based sampling offers a viable alternative. Using this approach, we demonstrated that our p16-associated signature was significantly enriched for BuckSenOvary, despite originating from independent donors. Although not all genes in BuckSenOvary showed directional concordance with our profile, this could likely be attributed to differences in donor populations and the inherent heterogeneity of senescence itself which varies based on factors such the inducing stress and time since induction [41, 42].

The relationship between senescence and cancer is complex, reflecting both antagonistic and permissive roles. Senescence acts as a potent tumor-suppressive mechanism by enforcing irreversible cell-cycle arrest in damaged or at-risk cells, thereby preventing malignant transformation. However, senescent cells also secrete a wide range of factors as part of the senescence-associated secretory phenotype (SASP), including cytokines, growth factors, and proteases, which can reshape the microenvironment in ways that may promote tumor initiation and progression [43]. In our study, enrichment analyses revealed that senescence-associated gene sets in ovarian cancer datasets exhibited directionally opposed regulation of many cell cycle–related genes, consistent with the growth-arresting function of senescence. In contrast, ECM-related genes demonstrated concordant changes across multiple cancer datasets, supporting the view that senescence-driven alterations to tissue structure and fibrosis may create a microenvironment permissive for tumor growth, although further investigation is warranted given that ovarian cancer can often originate in the fallopian tube and metastasize to areas such as the peritoneal cavity [16, 17, 38, 39].

Several limitations should be considered when interpreting the findings of this study. First, the analysis was conducted using tissue from only three postmenopausal donors, two of whom were of similar age and approximately two decades older than the third. The applicability of the senescent signature identified here needs to be further investigated across the broader postmenopausal population, including different stages of ovarian aging and health backgrounds.

Second, the proteomic analysis relied on a targeted 570-plex immuno-oncology panel. While this approach provided further insight beyond transcriptomics alone, it does not capture the full proteomic landscape. An untargeted and more comprehensive methodology, such as spatial mass spectrometry, would enable the detection of novel or unexpected protein changes associated with ovarian senescence [44].

Finally, the use of p16 as the defining marker for identifying senescence presents both strengths and limitations. p16 is a well-established marker of cellular senescence through its involvement in cell cycle arrest via inhibition of cyclin-dependent kinases [7-10], and the signature identified here supports senescence as a key process differentiating p16 positive and negative regions. However, p16 expression is not entirely specific to senescent cells; it can also be induced under certain stress or differentiation contexts, and not all senescent cells exhibit p16 upregulation [45]. This inherent heterogeneity is a common challenge in the senescence field, as no single marker can fully capture the complexity or diversity of senescent states across all cell types and tissues [22, 45].

Understanding the drivers and consequences of ovarian-specific senescence may have important implications for reproductive longevity and postmenopausal health. Interventions that modulate cellular senescence, such as senolytic or senomorphic compounds including rapamycin and quercetin, are currently being tested against reproductive aging, though they are in their infancy [46]. Hormone replacement therapy could also potentially alleviate ovarian senescent cell accumulation in postmenopausal women [47]. Here, the presence of p16-based differences despite all women being post-menopausal suggests that distinct rates of ovarian aging may occur, potentially shaped by genetic, environmental, or metabolic influences. Given that ovarian aging is tightly linked to the depletion of the primordial follicle reserve and the decline of estrogen production, determining whether senescence contributes to this decline or arises because of it represents a critical next step. Elucidating these relationships could reveal new methods for preserving ovarian function, mitigating age-related decline, and improving health outcomes in postmenopausal women.

This work provides understanding of the endogenous accumulation and spatial heterogeneity of senescent cells within the postmenopausal ovarian cortex. Future studies comparing these senescent profiles to those of younger, premenopausal ovaries will be essential to determine when and where senescence first emerges in the ovarian lifespan, and whether its spatial patterning influences tissue function, follicular depletion, and systemic health outcomes.

## Materials & Methods

### Histological staining and processing of ovary samples

De-identified human ovarian tissue was obtained from the Northwestern University Reproductive Tissue Library (NU-RTL) and processed and stained for p16 antibody using immunohistochemistry (IHC) followed by imaging on the EVOS FL Auto Cell Imaging System (ThermoFisher Scientific, Waltham, MA, USA) as previously described [15].

### Reference-guided TMA construction

Following quantification, these digitized images were used as a reference to identify 600 µm p16^INK4A^-positive and p16^INK4A^-negative areas of interest for each subject. These regions were then aligned with the corresponding donor paraffin blocks and sampled for transfer into a recipient tissue microarray (TMA) block, as previously described [48]. This reference-guided approach ensured that arrayed cores precisely corresponded to regions objectively defined during digital quantification.

### Sample preparation for spatial proteogenomics (SPG) on the GeoMx Digital Spatial Profiler

TMA sections (5 µm) were mounted onto Bond Plus slides (Leica Biosystems, Vista, USA) according to manufacturer recommendations. Sample preparation followed the NanoString SPG User Guide (MAN-10158) as previously described [49, 50].

Three antibodies were used as morphological markers for immunofluorescent staining: an anti-Transgelin antibody (clone SM22-alpha; Novus, catalog NBP3-121157) conjugated to Alexa Fluor 594 (1:100 dilution), a Pan-cytokeratin (PanCK) antibody (clone AE-1/E-3; Novus, catalog NBP2-33200) conjugated to Alexa Fluor 488 (1:400 dilution), and the nucleic acid stain SYTO-83 (Invitrogen, catalog S11364) conjugated to Cy3 (1:10,000 dilution). The Transgelin antibody was sheep-derived, and the PanCK antibody was mouse-derived.

### GeoMx Digital Spatial Profiling data acquisition

Digital spatial profiling was performed using the GeoMx Digital Spatial Profiler (NanoString Technologies, Seattle, USA) according to the manufacturer’s user guide (MAN-10152). FFPE slides were scanned across three fluorescence channels (FITC, Cy3, and Texas Red) to visualize morphology markers. Regions of interest (ROIs; 250 µm × 200 µm) were defined using GeoMx DSP software v2.0 and covered the full area of each profiled TMA core. Photocleaved oligonucleotide tags from each ROI were collected into individual wells of a 96-well PCR plate.

### Library preparation and sequencing

Library preparation was performed following the GeoMx DSP SPG protocol (MAN-10158). Libraries were sequenced by the University of Chicago sequencing core using an Illumina NovaSeq X platform in paired-end mode (Read 1: 38, index 1: 8, index 2: 8, read 2: 38), according to the manufacturer’s instructions.

### Initial processing of transcriptomic and proteomic data

Raw FASTQs were converted to digital count conversion (dcc) files with GeoMx NGS Pipeline (v.3.1.1.8). Segment-level sequence quality control was performed in R (v4.2.3) with GeoMxTools (v3.2.0), applying default thresholds for aligned (80%), trimmed (80%), stitched (80%), and sequencing saturation (50%) reads. Quantile-normalized counts were used for visualization of ROI clustering by uniform manifold approximation and projection (UMAP) with Seurat (v5.1.0).

Transcriptomic and proteomic counts were aggregated to the core level, filtered for low expression, and analyzed for paired differential expression and translation with EdgeR (v.3.40.2) using *voomlmFit()* and the model 0+∼P16, with subject included as a random effects block. Significance was called |log_2_FC| > 0.5 and *p*-value < 0.01. Correlation between the transcriptome and proteome was calculated using Spearman rank correlation. Only genes with unique transcript-to-protein mapping were included in correlation analysis (*n*=417). Genes subject to alternative splicing, post-translational modification, or shared protein products were excluded.

### Cell type deconvolution of transcriptomics

Cell type proportions within each segment were estimated using SpatialDecon (v1.8.0), with reference to an ovarian cortical single cell transcriptional dataset [18]. To generate the pseudobulk cell type signature matrix, cell types from a donor with less than 5 cells were removed, raw counts were normalized to counts per million (CPM) and averaged per cell type per donor. Genes were restricted to those within the GeoMx probes and top 200 markers per cell type (ranked by sign(log_2_FC) × −log_10_(*p*-value)). The median expression across donors within each cell type was then used to define the final cell profile matrix. Core-level cell type proportions were determined by averaging the percentage of each cell type across segments.

### Gene set overrepresentation analysis

clusterProfiler [51] was used to perform overrepresentation analysis of GO biological processes and KEGG pathways on the significantly upregulated and downregulated transcripts. A network illustrating the associations of genes and pathways was created with Cytoscape [52].

### Comparison with other senescence gene sets

Gene set enrichment analysis (GSEA) of the differentially expressed genes was carried out against the p16-associated ovarian signature, BuckSenOvary [15], a previously published doxorubicin-induced signature of ovary cortex explants [18] and other known senescence gene sets (GO Cellular Senescence (GO:0090398), Reactome Cellular Senescence (R-HSA-2559583), SenMayo [8], and the Senepy universal human signature [9]) using the fgsea package (v1.24.0) [53] in R.

### Comparison of differentially expressed genes with ovarian cancer datasets

To evaluate the enrichment of p16-associated significantly upregulated and downregulated gene sets in ovarian cancer, disease information of all available microarray- and RNA-seq based differentially expressed gene datasets in DiSignAtlas [37] was downloaded and datasets were filtered for organism=Homo sapiens, tissue=ovary, and diseases containing the words ‘cancer’, ‘tumor’, or ‘carcinoma’. Differentially expressed genes were ranked by sign(log_2_FC) × −log_10_(*p*-value). To reduce arbitrary ranking, genes with a log_2_FC, *p*-value, or adjusted *p*-value equal to N/A, a log_2_FC equal to 0, or a duplicated gene symbol were removed. Rankings were reversed when checking for enrichment of the p16-associated downregulated gene set. GSEA was then performed on the remaining datasets with fgsea.

